# TCR2HLA: calibrated inference of HLA genotypes from TCR repertoires enables identification of immunologically relevant metaclonotypes

**DOI:** 10.1101/2025.07.18.665436

**Authors:** Koshlan Mayer-Blackwell, Anastasia Minervina, Mikhail Pogorelyy, Puneet Rawat, Melanie R. Shapiro, Leeana D. Peters, Emily S. Ford, Amanda L. Posgai, Kasi Vengesana, Samuel Minot, David M. Koelle, Victor Greiff, Philip Bradley, Todd M. Brusko, Paul G. Thomas, Andrew Fiore-Gartland

**Affiliations:** Vaccine and Infectious Disease Division, Fred Hutchinson Cancer Center, Seattle, USA; Vaccine Immunology Statistical Center, Fred Hutchinson Cancer Center, Seattle, USA; St. Jude Children’s Research Hospital, Memphis, USA; Department of Immunology, University of Oslo and Oslo University Hospital, Oslo, Norway; Department of Pathology, Immunology, and Laboratory Medicine, Diabetes Institute, College of Medicine, University of Florida, Gainesville, FL, USA; Department of Medicine, University of Washington, Seattle, Washington, USA; Department of Global Health, University of Washington, Seattle, Washington, USA; Benaroya Research Institute, Seattle, Washington, USA; Department of Laboratory Medicine and Pathology, University of Washington, Seattle, Washington, USA; Imprint Labs, LLC. New York, NY, USA; Public Health Sciences Division, Fred Hutchinson Research Center, Seattle, USA; Department of Pediatrics, Diabetes Institute, College of Medicine, University of Florida, Gainesville, FL, USA; Department of Biochemistry and Molecular Biology, College of Medicine, University of Florida, Gainesville, FL, USA

## Abstract

T cell receptors (TCRs) recognize peptides presented by polymorphic human leukocyte antigen (HLA) molecules, but HLA genotype data are often missing from TCR repertoire sequencing studies. To address this, we developed TCR2HLA, an open-source tool that infers HLA genotypes from TCRβ repertoires. Expanding on work linking public TRBV-CDR3 sequences to HLA genotypes, we incorporated “quasi-public” metaclonotypes – composed of rarer TCRβ sequences with shared amino acid features – enriched by HLA genotypes. Using four TCRβseq datasets from 3,150 individuals, we applied TRBV gene partitioning and locality-sensitive hashing to identify ∼96,000 TCRβ features strongly associated with specific HLA alleles from 71M input TCRs. Binary HLA classifiers built with these features achieved high balanced accuracy (>0.9) across common HLA-A (9/12), B (9/12), C (6/13), DRB1 (11/11) alleles and prevalent DPA1/DPB1 (6/10), DQA1/DQB1 (8/17) heterodimers. We also introduced a high-sensitivity calibration to support predictions in samples with as few as 5,000 unique clonotypes. Calibrated predictions with confidence filtering improved reliability. Beyond genotype imputation, TCR2HLA enables the discovery of novel HLA- and exposure-associated TCRs, as shown by the identification of SARS-CoV-2 related TCRs in a large COVID-19 dataset lacking HLA data. TCR2HLA provides a scalable framework for bridging the gap between TCRseq data and HLA genotype for biomarker discovery.

## Introduction

Highly diverse αβ T cell receptors (TCRs) survey peptides presented by human leukocyte antigen (HLA) presenting molecules. Although TCR repertoires are routinely sequenced from blood and tissues, identifying immunologically relevant features within diverse adaptive immune receptor repertoire (AIRR) data remains a challenging problem [1,2]. Within each individual, highly polymorphic HLA molecules restrict the range of peptides available to interact with TCRs. Sufficiently large cohorts can be leveraged to identify the combinations of immunological exposure and HLA genotypes [3–5] that increase the likelihood of observing specific TCR features above the prevalence modulated by V(D)J recombination and thymic selection [3]. The capacity to evaluate TCR feature enrichment following immunological exposures across datasets in an HLA-defined manner would be particularly helpful in identifying relevant peptide-HLA epitope interactions that underpin T cell-mediated immunological control or pathology.

HLA alleles are encoded by distinct genetic loci. The class I loci – HLA-A, HLA-B, and HLA-C – encode proteins that complex with β2-microglobulin to present intracellular peptides to CD8⁺ T cells, while the class II loci – HLA-DRA1/DRB1, HLA-DQA1/DQB1, and HLA-DPA1/DPB1 – encode heterodimers that present peptides to CD4⁺ T cells [6–8]. To address the challenge that HLA genotypes are not universally collected in TCR repertoire sequencing (TCRseq) studies, recent work has explored the utility of using public TCR features to infer HLA genotypes from bulk TCRβ repertoires [9–11]. In this study, we exploit the fact that highly biochemically similar TCRs are often found across repertoires, such that searching for meta-clonotypes can increase feature publicity and reveal HLA-associated TCRs and motifs [11–13]. Due to biases in V(D)J recombination, many TCR sequences are uncommon and unlikely to be shared between individuals; however, rare non-identical receptors may undergo clonal expansion following antigen exposure [3,14–16]. Thus, we used near-exact neighborhood tabulation to capture “quasi-public” TCRs – rarer sequences [17] that, despite not being broadly shared themselves, occur within sequence neighborhoods that are public enough to support statistical association with specific HLA alleles. For inference of HLA genotype directly from repertoire data, we then developed models that combine (i) exact public features with (ii) sparsely public near-exact TCRs to build a binary classifier for each allele.

This study contributes open-source software that permits the discovery of exact and near-exact HLA-associated TCRβ features across thousands of input repertoires. The resulting high-confidence HLA genotype inferences allow for the identification of novel TCR:HLA associations in datasets lacking genotype labels. This facilitates the integration of labeled and unlabeled repertoires for discovering exposure- and HLA-associated TCRs that could serve as biomarkers for infectious disease, autoimmunity, cancer, and vaccine applications.

## Results

### TCR2HLA identifies exact and near-exact HLA-associated TCR features across cohorts

TCR2HLA is a statistical tool for inferring HLA genotypes from the occurrence patterns of receptors in peripheral blood TCRβ repertoires. Leveraging a training set of 3,125 repertoires from four independent cohorts with HLA-labeled data[18–21], we identified exact and near-exact TRBV-CDR3β features associated with common HLA alleles (**Fig. 1a-b**). Building on prior work identifying public TRBV-CDR3 sequences linked to HLA genotypes, we aimed to incorporate additional, “quasi-public” clonotypes with HLA-association evident based on detection of a TRBV-CDR3 AA feature or any single mutational variant. Notably, the probability of generation of TCRs with near-exact HLA-associated neighborhoods tended to be lower than that of exact TCR features (**Fig. S1a**). Combining distinct sets of exact and near-exact TRBV-CDR3β amino acid features (**Supporting Table S1**), we fit allele-specific classifiers via L1-regularised logistic regression, retaining the subset of features with non-zero weights (**Supporting Table S2**). Models used inverse statistical weighting to address class imbalance and post-estimation recalibration to standardize predicted probabilities across alleles and account for repertoire depth, helping to distinguish high-confidence positive or negative genotype calls from ambiguous ones (**Fig. 1c-d**). We evaluated model performance in multiple external validation cohorts with varied TCRseq amplification protocols and differing levels of richness (**Fig. S2, Supporting Table S3**) before we applied TCR2HLA models to 1,355 previously ungenotyped repertoires from SARS-CoV-2–exposed individuals, identifying novel exposure-associated meta-clonotypes through clustering of similar TCR sequences within HLA-associated sequence neighborhoods. More detailed descriptions of feature discovery, model training, and validation are provided in the Methods section.

**Figure 1:**
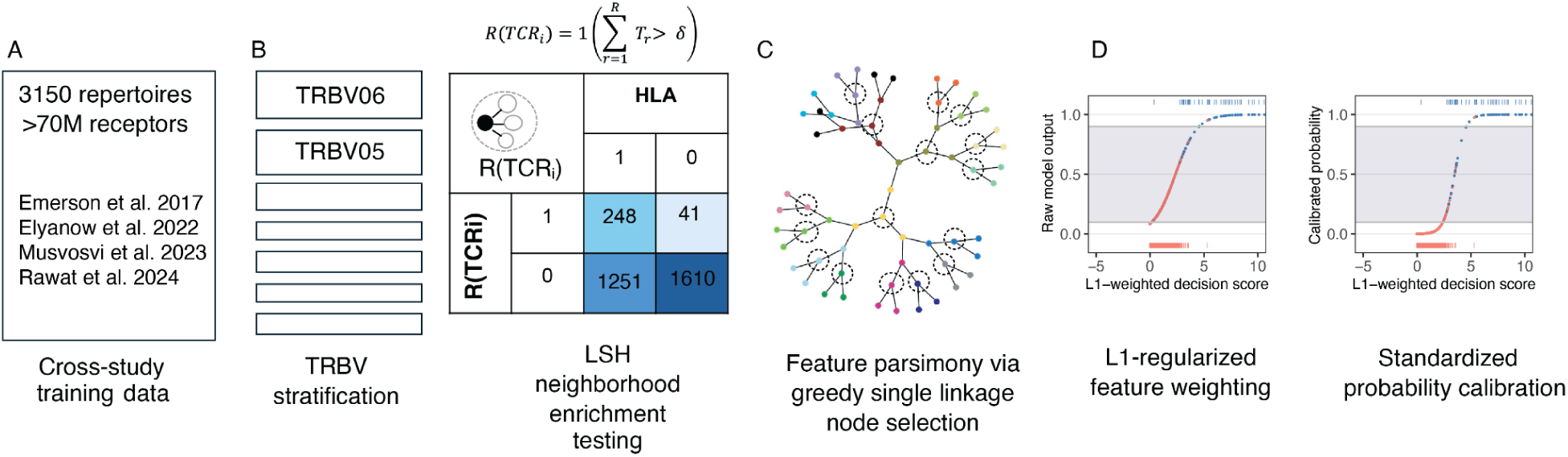
Schematic for generating calibrated HLA genotypes from TCRb repertoires. Schematic of statistical framework for discovering near-exact TCR features (defined by TRBV-family + CDR3 (amino acid)) in a multi-cohort wide association test. (**1a**) 71 M receptors present in >1 copy were assembled across 3150 repertoire from 4 studies. (**1b**) Use of TRBV-family stratification combined with locality-sensitive hashing facilitated a framework for discovery of exact and inexact HLA-associated TCR features. Fisher’s exact test is applied to an indicator function defined over the set {R} of neighboring TCRs to TCRi, where neighbors are defined as those TCRs within a mutational distance of 1 (i.e., differing by a single substitution, insertion, or deletion in the CDR3 sequence) and detected at a frequency exceeding the threshold ઠ=2×10^−6^. (**1c**) A greedy single linkage algorithm was used to reduce feature redundancy and potential variable collinearity prior to learning feature weights in L1-regularized logistic regression. (**1d**) Raw outputs from class-weighted L1-logistic regression optimized for ROC-AUC were recalibrated to standardized probabilities, enabling standardized decision thresholding and confidence-based sample retention for downstream analysis.

#### Allelic prevalence is similar across training and validation data sets

The prevalence of common HLA class I alleles was similar among the training data and the U.S. based validation datasets (**Fig. 2A**). By contrast, the prevalence of class II alleles differed between the training and validation sets. As expected, alleles associated with T1D risk, particularly DRB1*03:01, DQA1*03:02, DQA1*05:01, DQB1*03:02, and DQB1*02:01 and associated haplotypes, were enriched in the training data participants relative to those in the external validation datasets. Given strong geographical effects on the distribution of certain HLA alleles in human populations, we also examined the prevalence of HLA alleles from participants in a study conducted in South Africa (**Fig. 2A**, Musvosvi dataset). Reflective of the diversity represented in this dataset, certain alleles – particularly A*01:01, A*02:01, A*03:01, B*08:01, B*44:02, C*07:02 – were underrepresented in the South African cohort (**Fig. 2A**, **Supporting Table S4**). Some alleles (e.g., HLA-A*20:05, B*58:02, B*42:01, DRB1*15:03) that were strongly enriched in the South African participants but were not considered in the development of the cross-study common allele model because they were rare in the training and primary validation datasets.

**Figure 2:**
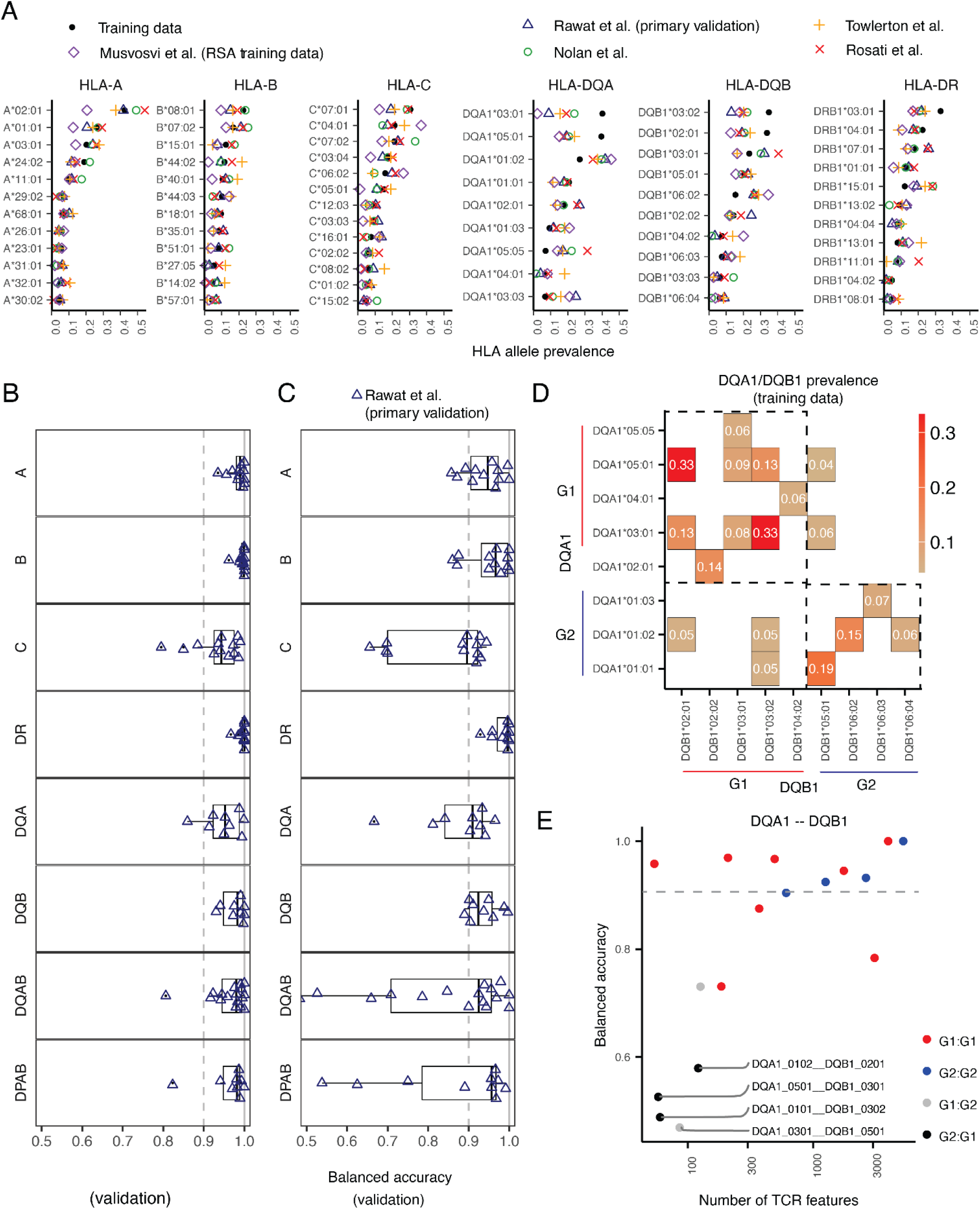
TCR2HLA classifier performance varies by locus on external validation datasets. (**2a**) Prevalence of alleles in training data versus validation datasets (Rawat et al., cohort 3 (primary validation), Towlerton et al., Nolan et al. and Rosati et al.). The Musvosvi et al. dataset represents a subset of the training data that includes participants from the Republic of South Africa (RSA). The model was fit for alleles or allele heterodimer (DQ, DP) combinations present in greater than four percent of overall training data – HLA-A (alleles, n = 12), HLA-B (n = 12), HLA-C (n = 13), HLA-DR (n = 11), HLA-DQA (n = 10), HLA-DQB (n = 10) HLA-DPA (n = 4), HLA-DPA (n = 6), and HLA-DQAB (n = 17) and HLA-DPAB (n = 9). Prevalence of HLA-DP, HLA-DPAB heterodimers, and HLA-DQAB heterodimers not shown (see **Supporting Table 4**) (**2b**) Receiver operator area under the curve (AUC) in the primary external validation cohort (adults 18-65, unique TCR > 25,000) by allele. (**2c**) Balanced accuracy (average of sensitivity and specificity) in the primary external validation cohort (adults 18-65, unique TCR > 25,000) by allele. (**2d**) Prevalence of DQA1/DQB1 group 1 (G1:G1), group 2 (G2:G2), and cross-group (G1:G2 or G2:G1) heterodimers in training data (**2e**) Balanced accuracy and number of unique TCR model features for each DQA1/DQB1 heterodimer. Colors indicate within-group or cross-group pairings.

### Common HLA genotypes can be inferred based on the presence of TCR features

To estimate model performance on healthy adult HLA-unlabeled TCRβ repertoires, we evaluated each common-allele model using validation repertoires that were neither used for feature discovery nor weighting. The model’s decision scores robustly separated samples by their HLA-status (AUC > 0.95) for most alleles across HLA-A, HLA-B, HLA-DR, and HLA-DQ loci (**Fig. 2b**). We also assessed the models’ balanced accuracies (BA) – measuring the ability to correctly predict positive and negative cases with a predefined decision threshold (**Fig. 2c**). In the primary validation repertoires, models correctly predicted the HLA-status with high balanced accuracy (BA > 0.9) for some, but not all alleles, with performance differing by HLA-locus: HLA-A (9/12 alleles), B (9/12 alleles), C (6/13 alleles), DRB1 (11/11 alleles), DQA1 (6/9 alleles), and DQB1 (8/10 alleles), as well as for DPA1/DPB1 (6/10) and DQA1/DQB1 (9/17) heterodimeric pairings (performance by allele, **Supporting Table S5**). Using the standard calibration (unadjusted for total repertoire diversity), models had high specificity on the primary validation data (i.e., median .98, IQR [0.95,0.99]) but a wider range of sensitivities across alleles (median 0.92, IQR [0.79,1]), with incorrect predictions across all models split between false positives (3.3%) and false negatives (2.2%) (**Supporting Table S6**).

After considering models for individual DQA1 or DQB1 alleles, we trained separate classifiers for DQA1/DQB1 heterodimers. While most models for common DQA1/DQB1 dimers had moderate to high balanced accuracy (>0.8-1.0 BA), the models predicting certain DQA1/DQB1 heterodimer pairings had low accuracy (BA < 0.6), despite the co-occurring alleles being prevalent in > 4% percent of training participants (**Fig. 2d**). At the DQ loci, two main classes of heterodimers are predicted to form stable HLA-DQαβ molecules: Group 1 (DQA1*02/04/05/06α+DQB1*02/03/04β) and Group 2 (DQA1*01α+DQB1*05/06β) [22,23]. Model accuracy for most Group 1 (G1α:G1β) and Group 2 (G2α:G2β) heterodimers was high. However, the remarkably low model accuracy for some heterodimers (e.g., DQA1*03:01/DQB1*05:01, DQA1*01:01/DQB1*03:02, DQA1*05:01/DQB1*03:01, **Fig. 2e**) is consistent with the hypothesized structural incompatibility of some cross-group pairings (i.e., G1α:G2β and G2α:G1β).

#### Model recalibration boosts balanced accuracy of genotype prediction in shallowly sequenced adult repertoires

The alpha-diversity of the primary validation repertoires (**Fig. 3a**) resembled that of repertoires used to train the model (**Fig. 3b**), however, other TCRseq datasets can have fewer unique TCR clonotypes per repertoire. To evaluate model performance in lower-diversity repertoires, we applied our HLA prediction model to data from Towlerton et al.[24], where many samples contained only 5,000–25,000 unique TCRβs – substantially fewer than in the primary validation set (25,000–450,000 unique TCRβs per sample; **Fig. 3a, 3c)**. Despite high class I HLA AUROCs (**Fig. 3e**), balanced accuracy was lower using the standard threshold calibrated from training data (P(HLA=1) > 0.5, **Fig. 3f**), due to fewer detections of HLA-enriched features in low diversity samples. The lower AUC and predictive performance observed for class II alleles in the Towlerton et al. validation dataset may be a consequence of depleted CD4^+^ T cell diversity expected in repertoires from participants living with human immunodeficiency virus (HIV), with one third of repertoires in that study collected prior to initiation of antiretroviral therapy[25].

**Figure 3:**
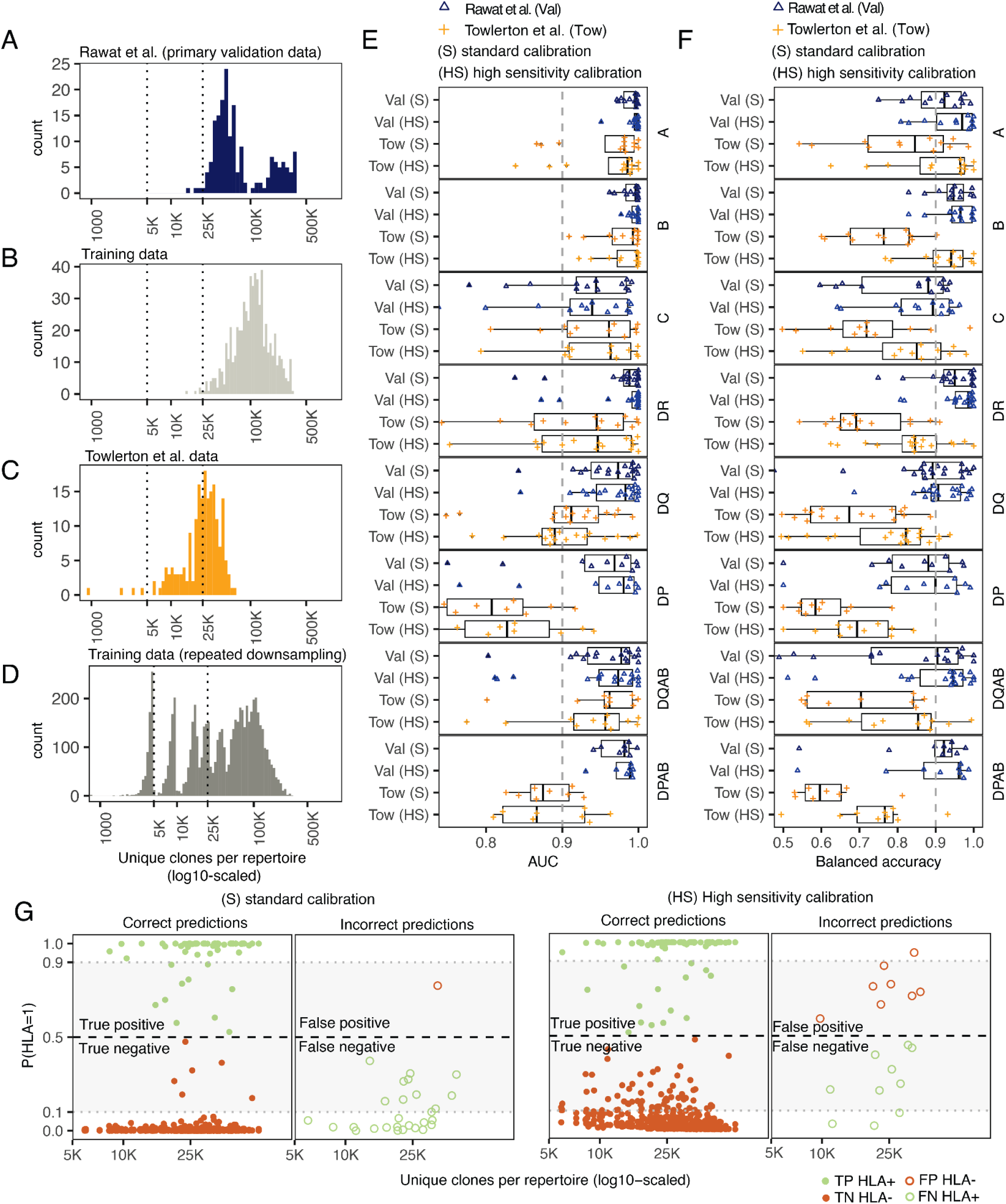
High sensitivity model calibration boosts sensitivity in lower diversity repertoires. (**3a,b,c,d**) Distribution of unique productive TCRb clonotypes sampled per repertoire in: (**3a**) Rawat et al. primary validation (Val, unique TCR > 5,000)) data (**3b**) training data, (**3c**) Towlerton et al. (Tow, unique TCR > 5,000) data with shallower depth of sampling (**3d**) and repeated downsampled training data. (**3e**) AUC in external validation datasets using either a standard or high sensitivity calibration. (**3f**) Balanced accuracy in validation datasets using either the standard (S) or high sensitivity (HS) calibration. (**3g**) Correct and incorrect predictions using the S or HS calibration. Prediction shown for A*03:01, A*02:01, A*01:01 alleles, where the colors of points show true HLA-status of the donor (green: HLA positive; orange: HLA negative), with closed dots indicating correct predictions (TP or TN) and open dots indicating incorrect predictions (FP or FN).

To improve HLA inference in lower diversity samples, we implemented a high-sensitivity (HS) recalibration, adding log10-transformed repertoire size to the model calibration with downsampled synthetic data (**Fig. 3d**, see Methods). For the Towlerton validation repertoires, the HS calibration improved balanced accuracy compared with standard calibration (IQR [0.72,0.91] vs. IQR [0.59,0.82] across all alleles) and reduced the number of false negatives, particularly in shallow repertoires (**Fig. 3g**) with a slight cost in specificity (IQR [.97-1] vs. IQR [.99-1] across all models). The HS calibration yielded more reliable probabilistic outputs, with fewer extreme false negative misclassifications. Overall, while recalibration markedly improved performance on shallow repertoires (Towlerton et al., median = 27,119 unique TCRβ), balanced accuracy achieved on these repertoires remained lower than in the primary validation set repertoires, possibly reflecting fundamental limits of inference from shallow sequencing data or in persons with highly depleted T cell memory. Further validation in additional DNA- and RNA-based TCRseq datasets showed sensitivity consistently dropping for repertoires with fewer than 25,000 unique clones (**Supporting Fig. S2, Supporting Table S5**).

#### Calibrated HLA genotype probabilities enable confidence-based label retention

While highly sensitive and accurate prediction was only possible for some HLA alleles, well-calibrated genotype probabilities can differentiate high from lower-confidence genotype inferences for those alleles that are more challenging to predict. To account for differences in model sensitivity across HLA loci and variability in repertoire sampling depth, we explored the use of calibrated HLA-genotype probabilities to implement confidence-based label retention. If probabilities are well-calibrated, masking of intermediate-confidence predictions should substantially increase balanced accuracy in retained data.

Thus, to assess the model calibration and feasibility of foregoing prediction in cases where the information content of a repertoire is low, we retained only high-confidence predictions (P < 0.1 or P > 0.9; **Fig. 4a–b**). This resulted in masking less than 10% of the predictions for most HLA-A, -B, and -DR alleles, and less than 20% of predictions for HLA-C and -DQ alleles (**Fig. 4c**). For HLA-DP alleles, this procedure resulted in data loss due to more intermediate-confidence predictions. After masking, median balanced accuracy exceeded 0.95 across most loci. While median accuracy among HLA-C alleles was less than 0.9 after data filtering, high balanced accuracy was possible for some HLA-C alleles (i.e., C*03:04, *04:01, *05:01, *06:02, *07:01). These results show that confidence-based masking can yield high-fidelity HLA labels for many alleles while retaining allele-level predictions from ≥ 80% of the repertoires.

**Figure 4:**
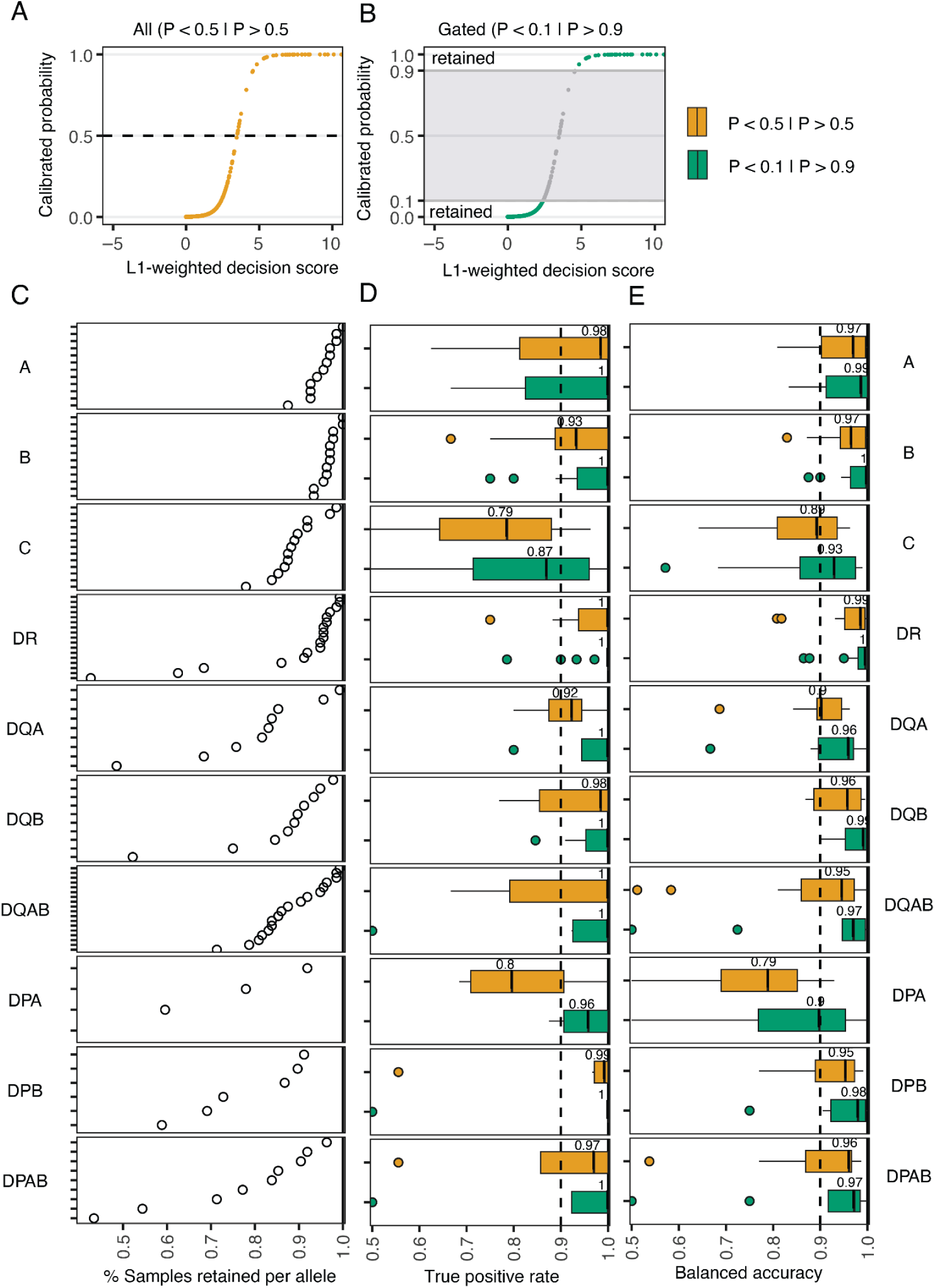
Confidence-based sample retention increases sensitivity and accuracy. (**4a**) Representative score reliability curve, showing mapping of classifier’s raw decision score to calibrated probabilities. When all data is considered (yellow) the prediction is based on a threshold probability of 0.5. (**4b**) Curve with intermediate probability estimates masked (0.1 > P > 0.9) to retain high-confidence negative and positive predictions. (**4c**) Fraction of samples retained per allele after confidence-based gating in the primary validation data not used for feature discovery or feature weighting from adults aged 18-65 with > 25,000 unique clones (n = 144). (**4d**) True positive rate of all (yellow) and high-confidence (green) predictions. (**4e**) Balanced accuracy for all (yellow) or high-confidence (green) predictions.

### HLA-inference facilitates the discovery of new HLA- and exposure-associated meta-clonotypes

Accurate HLA prediction is particularly valuable because it increases the possibility of discovering infection- or vaccination-associated TCRs in repertoires lacking explicit HLA genotype information. To this end, we applied TCR2HLA to 1,355 repertoires from individuals with acute or convalescent COVID-19 (Snyder et al.). We sought to use high-confidence genotype inferences (P < 0.1 and P > 0.9) to search the repertoires for novel HLA-associated receptors potentially associated with their common recent immunologic exposure. Using the inferred participant genotypes, we found numerous TCRs that were enriched in participants expressing specific alleles, but that did not match previously identified HLA-associated TCRs, including for representative allele DRB1*01:01 (**Fig. 5a**). A subset of the TCRs statistically associated with DRB1*01:01 formed distinct public sequence clusters (**Fig. 5b**). Because the reference datasets used to impute HLA labels were derived primarily from pre-2019 repertoires, we hypothesized that the unique TCR clusters discovered in the Snyder et al. cohort reflect expansions of SARS-CoV-2 reactive clonotypes.

**Figure 5.**
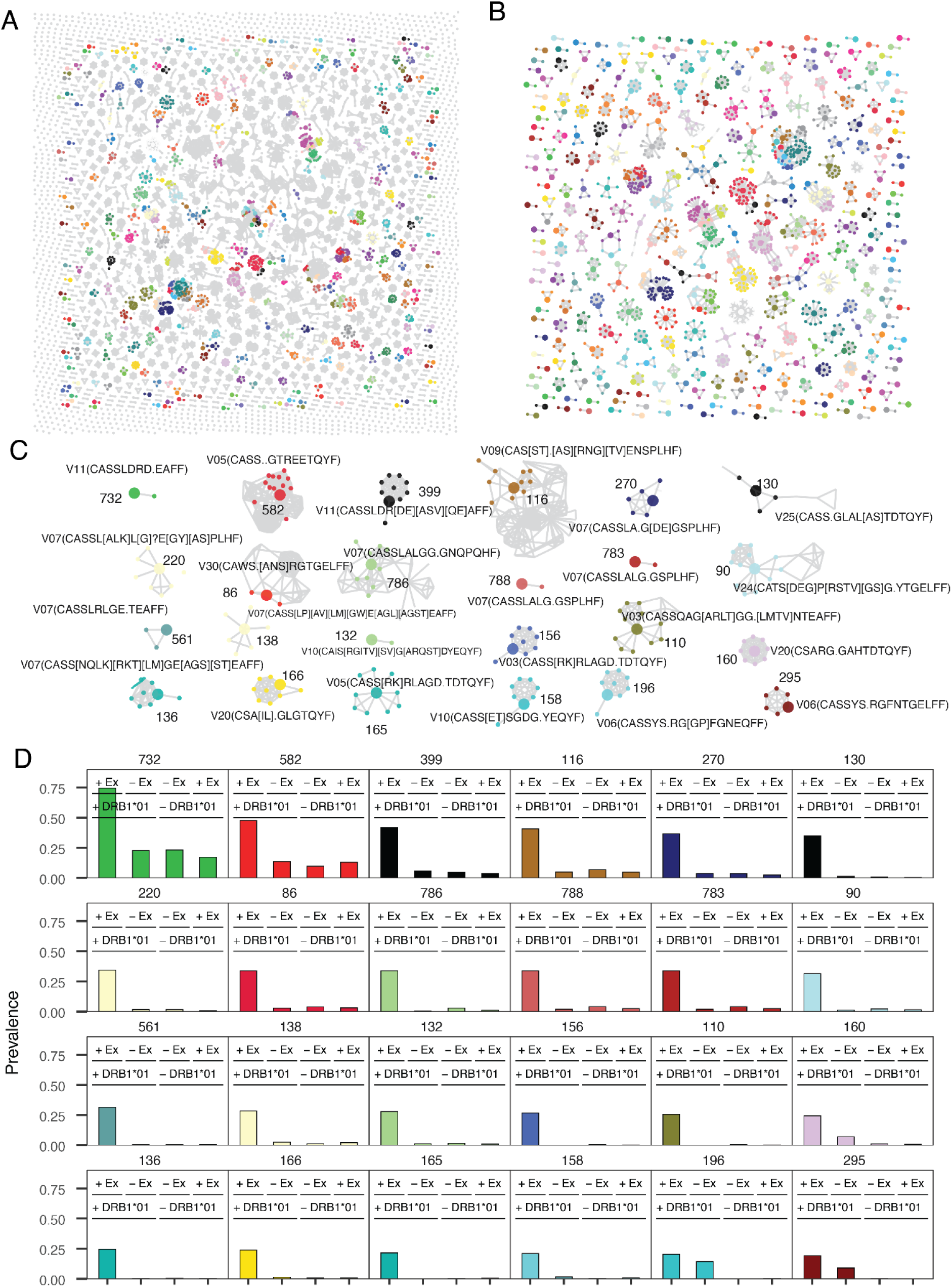
Identification of new HLA-associated TCRs in COVID-19-exposed repertoires. (**5a**) Sequence similarity network of TRBV-CDR3 clonotypes with DRB1*01:01-associated edit-1 neighborhoods. Gray nodes represent clones within one mutation of a previously identified DRB1*01:01 reference clone used for HLA genotype inference with TCR2HLA. Colored nodes indicate potentially novel DRB1*01:01-associated TCRs that may reflect recent SARS-CoV-2 exposure in this cohort (n = 1,355; Snyder et al., 2025). (**5b**) HLA-associated TCRs were clustered by sequence similarity (TCRdist ≤ 14) and grouped into meta-clonotypes using a degree-based greedy graph-based algorithm. Large nodes represent anchor clones with the highest number of first-degree neighbors. (**5c**) Twenty-four strongly HLA-associated meta-clonotypes, defined by CDR3 regular expression (regex) motifs and TRBV gene usage. Regex patterns were generated by aligning first-degree neighbors (smaller nodes) to their corresponding anchor TCRs (large nodes). (**5d**) Relative prevalence of meta-clonotypes in exposed (+Ex, n = 1,355) and unexposed (–Ex, n = 1,977) repertoires, stratified by DRB1*01:01 genotype (+DRB1*01). Colors match the meta-clonotypes identified in previous panels.

To reduce redundancy and engineer features generalizable to closely related receptors, we organized novel enriched TCRs into meta-clonotypes defined by an anchor sequence and its first-degree neighbors based on TCRdist-based similarity networks. These clusters, represented by a V-gene family and CDR3 regular expression, capture quasi-public TCRs hypothesized to recognize the same HLA-presented epitope (**Fig. 5c**). We screened the meta-clonotypes for their prevalence in individuals with and without the associated HLA allele in the Snyder et al. cohort and a pre-COVID cohort (Rawat et al. cohorts 1 and 2)[18,26]. When stratified by exposure and genotype, many of these novel HLA-associated meta-clonotypes showed specific enrichment in repertoires from SARS-CoV-2-infected individuals carrying the relevant restricting HLA allele (**Fig. 5d**), supporting their potential application as antigen-specific biomarkers.

A major strength of TCR2HLA and our cross-study reference set of broadly HLA-associated TCR clones is the potential to discover novel HLA-associated TCRs that are selectively enriched by exposure and HLA genotype. In this example, we show how TCR2HLA can be used to infer HLA status of a large set of unlabeled repertoires and identify new clusters of TCRs with strongly HLA-associated neighborhoods. This is particularly valuable for the identification of exposure-associated CD4^+^ T cell receptors, since compared to CD8^+^ T cells, this population is less identifiable by longitudinal expansion analysis, tetramer staining, or activation-induced markers, with potential confoundment by bystander activation [27,28].

### TCR2HLA models underperform when applied to TCR repertoires from children

The human TCR repertoire size and diversity change over the lifespan, in part, due to declining thymic output and clonal expansion of memory T cells [29,30], with diversity negatively associated with age and cytomegalovirus (CMV) exposure[31]. Thus, despite the utility of the TCR2HLA framework, we conducted sensitivity analyses to assess whether model performance varied by age. We hypothesized that predicting HLA status might be more difficult in children than adults due to higher repertoire diversity maintained by higher thymic output, fewer total exposures, and smaller T cell memory fractions in early life [32,33]. To test this, we evaluated classifier performance in childhood repertoires not used in training from the DAISY (Diabetes Autoimmunity Study in the Young) cohort described in studies by Mitchell et al. (2022, 2023), which included children aged 1–18, many diagnosed or at risk for type 1 diabetes [34,35]. Compared to observed performance in adults, balanced accuracy and AUCs were notably lower, even for common HLA alleles (representative results for HLA-B and HLA-DR, **Fig. 6**). While models retained high specificity, low sensitivity led to significantly lower accuracy in children than adults (p-value < 0.0001), which was lowest in children under 5 years and increased with age. A recalibrated model, using the original TCR features and decision scores, but calibrated only with training data from participants under 13 years (U13 model), did not significantly improve childhood predictions relative to calibration with the full training dataset (p-value = 0.3). Adolescents (ages 13–18) showed intermediate accuracy compared to young children, suggesting key HLA-linked TCR features may arise due to antigen exposures during adolescence or early adulthood. Reduced overall TCR diversity in adult repertoires may also enhance feature detection, as memory clones become enriched relative to the naive T cell population. We also noted a trend of slightly reduced accuracy for HLA-B alleles in adults over 65 years, despite consistently high AUCs, potentially reflecting immunosenescence or calibration biases, as most of the training data came from individuals under age 65.

**Figure 6:**
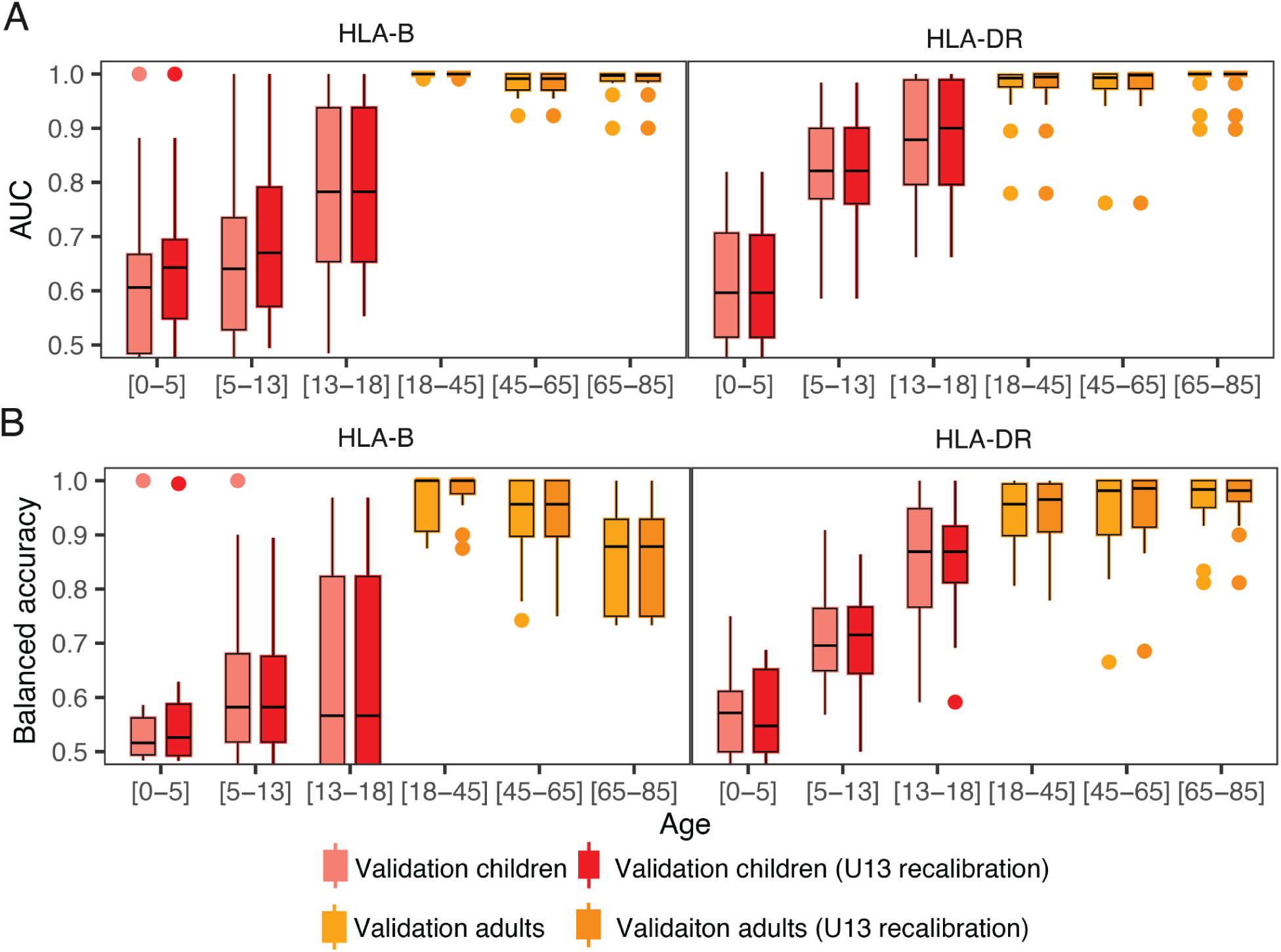
Sensitivity of HLA inference differs by age group. (**6a, 6b**) For selected performance and separating HLA-genotype of sample (AUC, **6a**) and predicting HLA-genotype (balanced accuracy, **6b**). Probabilities were calibrated either using full training data or only data from participants under 13 years of age (U13). External validation dataset from children longitudinally sampled, Mitchell et al., 2022) and standard validation dataset (n = 142, ages 18-65 years, and n = 32 > 65 years). Age category [0-5], [5–13], [13–18] includes children under the age of 5, 13, and 18, respectively.

### Scope and accuracy of HLA genotype inference are limited by a lack of ethnic and geographic diversity

Because HLA allele frequencies and immunological exposures vary substantially by geographic region, we evaluated the limitations of using a primarily U.S.-based training dataset to predict HLA status of individuals from the Republic of South Africa (RSA), who differ partially in their HLA genotype diversity and environmental exposures (e.g., higher prevalence of HIV and tuberculosis [36]). To assess this, we removed all South African repertoires (n = 144) from feature discovery, selection, model weighting, and calibration steps, generating a U.S.-only model (xRSA). We then compared the predictions (**Fig S3a**) and predictive accuracy (**Fig S3b-c**) of the full model (trained on all data) and the xRSA model on the RSA repertoires. We observed that for several alleles common to both U.S. and RSA cohorts (e.g., A*01:01 and A*02:01), the xRSA model retained similar accuracy. While this comparison is imperfect, as models generally perform better against data used in training, we did observe that prediction accuracy and AUCs were reduced after removal of RSA. training data for several common alleles, including A*11:01, A*24:02, B*44:03, C*07:01, C*07:02, C*16:01, DQA1*05:05, and DQB1*04:02 (**Fig. S3b-c**). However, the fact that the xRSA models’ AUROCs show that raw decision scores still robustly separate HLA genotype-positive and -negative samples among RSA participants, suggesting that many learned features are shared across geographic contexts. The reliability of the xRSA model declined relative to the full model, with more HLA-positive samples assigned moderate confidence scores (0.4 < P(HLA=1) < 0.9). In some models, such as DQA1*05:01, excluding RSA repertoires from model training led to an increased number of false positives, possibly because the xRSA model failed to downweight TCRs that, in the RSA context, are associated with a different HLA allele not as commonly found in the U.S. population.

Despite reasonable performance for many common shared alleles, it is important to recognize that many HLA alleles common among the RSA participants were extremely rare or altogether absent in the U.S. training data (e.g., representative HLA-B alleles in **Fig. S3d**; HLA-DR in **Fig. S3e**). For alleles present in 1–4% of the combined U.S.–R.S.A. dataset, we developed a rare-alleles model (**Supporting Table S2**), but we excluded from consideration models for alleles with <1% prevalence. Models trained primarily on U.S. data provided less coverage of the full allelic diversity among South African participants. For example, the common-allele model set developed from training data in this study included at least one HLA-A, -B, and -C allele for 95%, 87%, and 96% of U.S. participants, respectively – but only 76%, 54%, and 42% of R.S.A. participants (**Fig. 7a**). Including rare allele models (which could not be externally validated) expanded this coverage (at least 1 allele per loci) to > 89% of participants in both U.S. and R.S.A. datasets across considered HLA-class I and HLA-class II loci (**Fig. 7b**).

**Figure 7:**
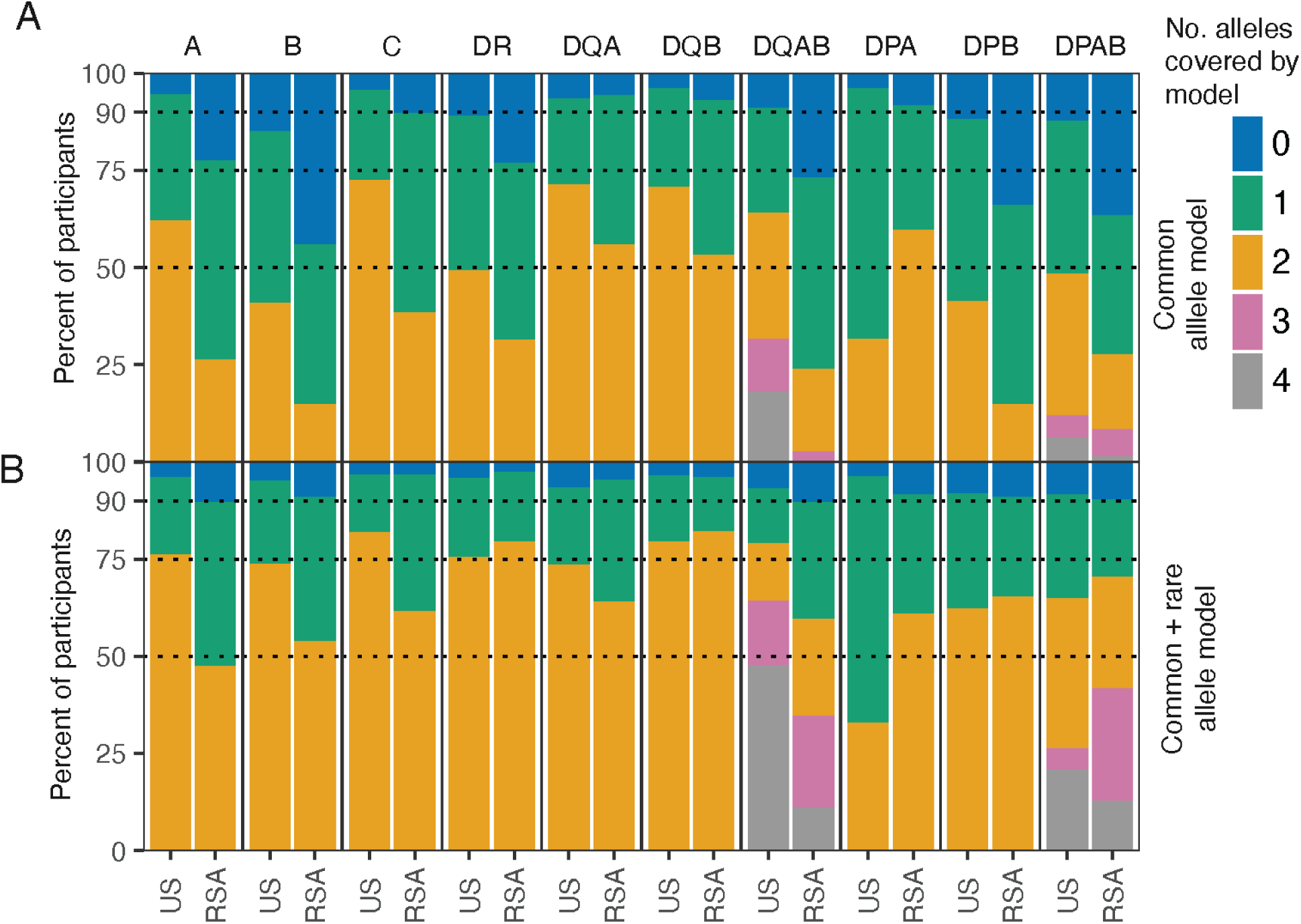
Model coverage of HLA-diversity differs by geographic region. (**7a**) Common allele model coverage of alleles by locus in training data from the United States (US) and the Republic of South Africa (RSA). (**7b**) Enhanced coverage by combined common and rare allele models.

## Discussion

In this study, we developed open-source software for the large-scale discovery and selection of HLA-associated TCR features. Our work builds on prior efforts to infer HLA genotypes from unlabeled TCR repertoire data. For example, Mayer-Blackwell et al. (2021) employed public HLA-associated TCRβ sequences previously identified by DeWitt et al. (2018) to predict expression of common HLA alleles in unlabeled repertoires. The inferred HLA status of each repertoire was used to evaluate TCR meta-clonotypes and GLIPH motifs – learned from MIRA assays – for evidence of HLA restriction after SARS-CoV-2 exposure [4,11,15,37]. Ortega et al. (2025) used L1-regularized models trained on TCRα and TCRβ features derived from Rosati et al. (2022), Russell (2022), and Emerson et al. (2017) datasets to predict participant HLA status [9,19,38,39]. Zahid et al. (2024, preprint) identified ∼1 million HLA-associated public TCR clones from 4,144 genotyped individuals but have not released those features or an open-source predictive tool [40]. Recently, ElAbd et al. (2025, preprint) developed an HLA imputation framework using public HLA-associated features from 5,554 TCRβ repertoires across cohorts from Germany, the U.S., and Norway, primarily involving individuals with Crohn’s Disease and Ulcerative Colitis [10]. Pan et al. (2025, preprint) applied the GIANNA clustering algorithm [41,42] to publicly available HLA-genotyped datasets, identifying CDR3s from GIANNA clusters predictive of HLA genotype. While these efforts all make unique contributions, our study offers a scalable method for discovering HLA-association among related sets of similar TCRs across studies and demonstrates how well-calibrated HLA-genotype probabilities estimated from the occurrence of non-exact TCR features can be leveraged to discover new HLA-restricted and exposure-associated receptors.

In this study, we integrated more than 1,900 HLA genotypes and TCRβ repertoires from a cohort comprising individuals with T1D, their close relatives without diabetes, and unrelated controls without diabetes [18] with 1,250 TCRβ repertoires from three other previously available TCRβ datasets to develop an open-source software for calibrated HLA genotype predictions. Expanding on Ortega et al.’s approach [9], we evaluated public feature tabulation strategies that use edit-1 neighborhoods instead of exact sequence matches. We first observed that “fuzzy” edit-1 tabulation reduced HLA specificity for many individual public HLA-enriched features with high probability of generation (TCRβ formed from commonly used TRBV- and TRBJ-genes with CDR3 junctions more easily formed by V(D)J recombination mechanism). Ortega et al.’s use of L1-penalized regression likely counteracted this through strong L1-penalized regression, effectively dropping fuzzy features lacking an HLA-specific signal. In contrast, in the present study, we fundamentally changed the way HLA-association testing was performed: rather than limiting enrichment tests to the neighborhoods of known public features, we extended *edit-1* tabulation into the feature discovery phase itself, testing millions of candidate neighborhoods across the multi-study cohorts. This allowed us to identify quasi-public TCRs with lower generation probabilities and robust HLA-specific enrichment that would be undetectable using exact-match methods alone.

To efficiently query neighborhoods among the ∼70 million TCRs in our input datasets, we employed TRBV-family data partitioning and locality-sensitive hashing [43] to rapidly detect highly similar TRBV-CDR3β sequences across thousands of repertoires. This approach parallels the GIANNA clustering strategy used by Pan et al. 2025 (preprint) [42], where vectorized CDR3 sequence embeddings were utilized for TCR clustering and HLA-enrichment testing. However, while the GIANNA algorithm initially clusters receptors using CDR3 information, the authors report that incorporating TRBV-gene information post-hoc boosted the HLA-signal [42]. In our study, we opted to define features at the TRBV-gene family level during the feature discovery phase to minimize batch artifacts that can arise at the TRBV-gene or allele level when combining datasets generated over time due to differences in amplification chemistries and advances in gene-calling pipelines.

A key aspect of our study was the evaluation of generalizability across heterogeneous datasets not used in feature discovery or model training, including repertoires varying from 5,000 to 450,000 unique clones. By calibrating prediction to stratify high- vs. low-confidence HLA genotype calls, we enabled efficient use of variable quality repertoires for downstream discovery of disease- or vaccine-specific HLA:TCR associations. Notably, the reduced sensitivity of *TCR2HLA* (high false negative rate) we observed in pediatric samples supports the hypothesis that enrichment of many public TCR-CDR3 features and neighborhoods results from common viral exposures and subsequent clonal expansion, rather than from imprinting by HLA proteins during thymic selection. This dependence on robust, exposure-driven response implies that TCR2HLA’s performance may decline in immunocompromised individuals or those with markedly different exposure histories.

We focused on inference of HLA genotypes from TCRβ repertoires, because TCRβ is the most commonly collected data type; however, the same approaches can be adapted to include TCRα occurrence alongside TCRβ data as was shown by Ortega and colleagues [9]. The advent of new technologies for generating deep snapshots of TCRαβ pairings [44] could be combined with single-chain HLA inference models to rapidly identify candidate TCRαβ sequences’ cognate HLA alleles to clone into cell lines for direct experimental validation. Future work might also take advantage of more sophisticated distance metrics to define a robust library of paired chain anchor features that could enhance HLA genotype inference beyond what is possible with single chain data alone or reveal TCRβ features that are conditionally linked to distinct HLA-alleles depending on α:β pairings.

As with other methods, *TCR2HLA* showed variable sensitivity across loci and by allele, motivating us to differentiate between high- and low-confidence predictions. Even when limiting to high-confidence imputations, a core limitation of the *TCR2HLA* tool is that it only currently predicts presence and absence of HLA alleles with > 4% frequency in the U.S.-based training data. As such, *TCR2HLA* makes predictions for only a fraction of global HLA allelic variation. In sensitivity analyses, model performance on South African repertoires decreased for some alleles when these repertoires were withheld from the feature discovery and weighting steps, underscoring potential limitations to transporting models to ethnically divergent cohorts. For some HLA alleles common in both populations (e.g., A*02:01, B*07:02, DRB1*07:01), removal of South African repertoires did not substantially diminish predictive performance, most likely because many common human viruses circulate globally with associated TCR responses shared across geographically diverse populations.

Although Epstein-Barr virus (EBV)-, CMV-, and influenza-related TCRs appear in geographically distributed populations, HLA diversity could theoretically impact immunodominance hierarchies and alter population-specific TCR occurrence patterns even for globally common alleles. As with peptide-HLA binding prediction tools [45,46], limited allele representation for low- and medium-income countries in training data remains a barrier to model generalization [47]. Given the diagnostic value of HLA genotyping for maximizing insight from TCRseq data, expanding the geographical and genetic diversity of reference HLA-labeled TCR datasets is essential. The framework and software developed in this paper provide a scalable path forward for extending HLA genotype inference and feature discovery to new global datasets as they become available, as well as identifying novel exposure-elicited HLA-associated TCR features that could be leveraged as infectious disease, cancer, or autoimmune biomarkers.

## Methods

### HLA-labeled training data

To develop predictive models of HLA genotypes from peripheral blood TCRβ repertoires, we assembled a training dataset of 3,125 repertoires from four studies (**Supporting Table 1**) and constructed a bioinformatic pipeline to identify exact and near-exact TCR features strongly associated with HLA alleles **(Fig. 1a,b,c**). The training data primarily included participants living in the United States from prior Emerson et al. (n = 555), Elyanow et al. (n = 276), and Rawat et al. (n = 1,977) studies [18,19,21]. We further included a study with participants recruited in South Africa (n = 144, repertoires = 317) [20]. The largest study in the training set was conducted to study risk factors associated with type 1 diabetes (T1D; n = 1,977); it includes individuals diagnosed with T1D as well as first-degree and second-degree relatives without T1D, and unrelated controls without T1D. Because certain class II alleles are strongly associated with the risk of developing T1D [48–50], the allelic prevalence in the training data is partially skewed from that of the general population of the United States and from other cohorts in the training data [51].

HLA alleles were inferred from genetic data at the 4-digit resolution. The methods for HLA-genotyping varied and are described in detail by study [4,18–21]. Briefly, participants in Rawat et al. were genotyped using the UFDIchip custom microarray, including more than 9000 markers covering the HLA genomic loci; genotypes were imputed from single nucleotide polymorphisms (SNPs) using Axiom™ HLA Analysis software (v1.2.0.38) [18]. Genotype data from Emerson et al. involved assembling genotypes from various methods as described by Dewit et al. [4]. From the Emerson dataset, we restricted our analysis to 555 repertoires from participants with genotyping coverage across common class I and class II HLA alleles. High resolution HLA genotyping for HLA class I and class II loci for participants in Elyanow et al. and Towlerton et al. studies was performed using Scisco Genetics two-stage PCR protocol – ScisGO HLA Typing kit v6 – designed for next-generation sequencing (NGS) [21,24,52]. Four-digit allele imputation accuracy of HLA-alleles from microarray-based SNP data varies by allele [53]; thus, we anticipate integrating this data with high-resolution NGS-derived HLA-genotype labels would improve the robustness of TCRβ feature discovery.

### Exact and near-exact HLA-associated feature discovery

Combining data across studies and HLA-genotyping methods, we attempted to identify robust cross-study HLA-associated TCR features, defined by TRBV family and CDR3β amino acid (AA) sequence (TRBV-CDR3), based on statistical enrichment in the repertoires of participants with specific HLA alleles (4-digit resolution). HLA inference was framed as a series of separate allele-level binary classification tasks (e.g., HLA-A*02:01 positive vs. negative). To limit the feature space, we enriched for expanded T cells, using only TRBV-CDR3AA features observed in >1 template per repertoire. Sequences were partitioned by IMGT V-gene family. Edit-distance-1 (one substitution, insertion, or deletion) neighbors within V-family partitions were identified via locality-sensitive hashing – a computationally efficient method of sequence similarity search relying on an index of position-masked strings [43] related to other index-based methods for identifying neighboring CDR3 sequences [15,54]. TCR features found in ≥5 training set repertoires were tested for association with each common HLA-allele by Fisher’s Exact Test (*fishersapi*), with strong associations defined by (edit-1, odds ratio > 1, p < 10^-8^, and absent in >90% HLA-mismatched repertoires).

For TRBV-CDR3 features with a strong HLA-association based on edit-1 detections, we constructed sequence similarity graphs with edges connecting observed TCRs within TCRdist < 14 (*tcrdist3* and *networkx*) [11,14]. To remove potentially spurious features (e.g., TCRs with HLA-enriched neighborhoods, but no individual HLA association), we pruned nodes that lacked any enrichment on an exact sequence basis (odds ratio < 1). To further reduce redundancy among remaining HLA-associated features, we applied a greedy algorithm to select “anchor” TCRs to retain for downstream prediction tasks: within each sequence similarity graph, the iterative algorithm selected the highest-degree node as a TCR anchor, then removed it and its neighbors from the graph, before iteratively selecting the next most densely connected anchor, until no nodes of degree > 1 remained. We also identified exact TRBV-family+CDR3 features: public HLA-associated features found across the four studies (p < 10^-8^, odds ratio > 1) as well as those previously identified from the Emerson et al. cohort (p< 10^-6^) as previously reported by Dewitt et al. [4].

### Feature selection weighting

Both highly public TCRs and sparsely public TCRs (with sufficiently public sequence neighborhoods to power an association) could be combined to build a binary classifier for each allele (**Fig. S1a**). We trained binary classifiers combining exact and edit-1 anchor features, estimating weights with binomial generalized linear models (GLMs) and L1 regularization. Five-fold cross-validation was used to tune regularization to achieve maximal area under the receiver operating characteristic curve (AUROC). Across all alleles, both edit-0 and edit-1 features were retained and assigned positive feature weights (**Fig. S1b**). There was a modest positive correlation between the strength of the statistical HLA-association in the feature discovery phase and the GLM-derived model weights (**Fig. S1b**). To address severe class imbalance, we used inverse class probability weighting in the model fitting stage, and post-estimation calibration by Platt’s method to refine predicted probabilities (**Fig. 1d**). For application to samples with low numbers of unique clonotypes (5-25K), we implemented a high-sensitivity (HS) recalibration, using a second-degree polynomial adjustment based on the number of unique clones (log_10_-transformed) with downsampled synthetic data.

For each allele, after selecting highly HLA-enriched TCRs across four cohorts, we trained models one allele at a time, based on a set of *n* informative exact TCRβ and a set of *m* near-exact anchor TCRβ features. Each sample is thus represented as a vector x ∈ {0, 1}^n+m^ composed of two binary subvectors:

- **x_0_** ∈ {0, 1}^n^: indicates if a sample contains at least one clonotype at productive frequency > 2E^-6^ matching that HLA-associated exact feature.
- **x_1_** ∈ {0, 1}^m^: indicates if a sample contains at least one clonotype at productive frequency > 2E^-6^ within one mutation (substitution, indel, or deletion) of an HLA-associated near-exact anchor feature

Such that:

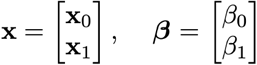

The probability that a repertoire comes from a person expressing an allelic variant (y) is expressed as:

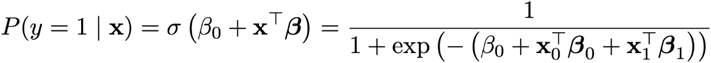

We used regularization to estimate a parsimonious set of parameters that maximizes the L1-penalized log-likelihood.

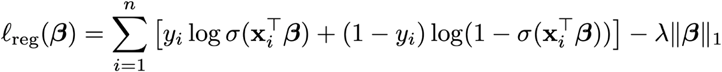

To account for extreme class imbalance, important for predicting the presence of rarer alleles, inverse probability weighting (IPW) was applied during model fitting.

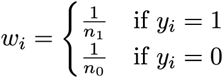

Where n_1_ and n_0_ are the number of allele-positive and allele-negative repertoires in the training data, respectively. The IPW-weighted L1-regularized log-likelihood is:

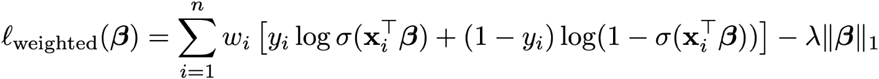

This objective function ensures that both classes contribute during training, even under severe class imbalance. The L1-regularized logistic regression with feature weights β and regularization parameter λ learned by maximizing AUROC across 5-fold cross-validation, yields a decision score.

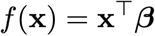

However, after fitting with inverse probability weighting and regularization, the decision score may not be well-calibrated as a probability. Thus, Platt scaling fits a sigmoid to the raw decision function:

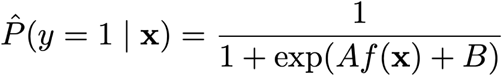

For this standard calibration, parameters A and B were then fit on the full training set. For the

high-sensitivity calibration – to account for low sample diversity < 25,000 unique clones – we include an additional adjustment using the covariate u_i_ = log_10_(U_i_), where U_i_ is the number of unique clonotypes (with frequency > 2E-6) in sample *i*. The calibrated probability is thus derived by adjusting the decision score as follows:

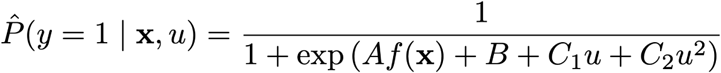

Where the parameters C_1_ and C_2_ are fit simultaneously with A and B on the repeatedly downsampled draws from the training set, with the decision function extended to include the first and second degree polynomial terms. The synthetic training data was generated from 650 training data repertoires repeatedly sampled with replacement for 1, 5, 10, 25, 50, 100, and 500 thousand draws.

### Assessing generalizable model performance via external validation

We tested model generalizability using several validation cohorts that were not used for feature discovery or model calibration. The primary validation, cohort 3 in Rawat et al., included 142 adults (18–65 years) unrelated to participants in the T1D cohorts 1,2 with >25,000 unique TCRs per sample.[18] We performed additional validation using a smaller COVID-19 cohort (Nolan et al., 63 repertoires), a dataset including low-diversity repertoires in HIV positive individuals (Towlerton et al., 188 repertoires), and an mRNA-based 5’-race derived TCRseq (Rosati et al., 112 repertoires)[18,24,26,38].

Models were built for common alleles (>4% prevalence in training data; **Fig. 2A, Supporting Table S2,** model v4) to ensure sufficient training and validation data were available to robustly examine allele and locus-specific predictive performance: HLA-A (n = 12 alleles), HLA-B (n = 12), HLA-C (n = 13), HLA-DR (n = 11), HLA-DQA (n = 10), HLA-DQB (n = 10), HLA-DPA (n = 4), HLA-DPB (n = 6). Separately, we also trained models for rare alleles (1-4% prevalence, **Supporting Table S2,** model v7), though evaluation of these models’ performance was limited due to insufficient validation data: HLA-A (n = 7 alleles), HLA-B (n = 21), HLA-C (n = 6), HLA-DR (n = 19), HLA-DQA (n = 2), HLA-DQB (n = 5), HLA-DPA (n = 1), HLA-DPB (n = 11). Due to inconsistent occurrence and genotype availability, we did not focus on TCRs related to DRB3, DRB4, or DRB5 loci.

### Meta-clonotype discovery from unlabelled repertoires

We applied TCR2HLA to infer HLA genotypes for 1,355 previously unlabeled TCR repertoires from individuals with acute and convalescent COVID-19 or high-risk exposure [26]. For common HLA alleles, genotype predictions were retained when confidence was high (calibrated probability > 0.9 for presence or < 0.1 for absence). We then used TRBV-family partitioning and locality-sensitive hashing to identify TCRs that shared HLA-associated edit-1 neighborhoods in the COVID-19 cohort. To identify potentially novel, cohort-specific TCRs, we excluded receptors highly similar to the reference TCRs used to make HLA inferences, defined as sharing the same TRBV family and differing by no more than one AA substitution, insertion, or deletion in the CDR3 region.

To reduce feature redundancy, we grouped newly discovered HLA-associated TCRs into meta-clonotypes based on strong statistical associations with HLA alleles (OR > 1, p < 1e-8), after excluding TCRs from individuals lacking the relevant HLA genotype. For each allele, we constructed sequence similarity networks by connecting TCRβ sequences with pairwise TCRdist ≤ 14 (typically a strict similarity arising from matching TRBV gene-usage and 1-2 CDR3 amino acid substitutions). A degree-based greedy algorithm was used to identify anchor TCRs – clones with the most first-degree neighbors – and define meta-clonotypes as anchor-neighbor clusters. Each meta-clonotype was defined by a regular expression (regex) motif derived from aligned CDR3 sequences [11,55] and annotated with the TRBV family of the anchor clone. We then quantified the occurrence of TCRs matching each meta-clonotype definition across the Rawat et al. (pre-COVID) and Snyder et al.(COVID-exposed) cohorts to evaluate prevalence by HLA genotype and exposure status [18,26].

### Data availability

TCR repertoires data used to identify HLA-associated features were analyzed from previous studies and are available (https://clients.adaptivebiotech.com/pub/emerson-2017-natgen, https://clients.adaptivebiotech.com/pub/elyanow-2022-jci, https://clients.adaptivebiotech.com/pub/musvosvi-2022-nm), and Rawat et al. data will be available for download from the AIRR Data Commons using the iReceptor Gateway using Study ID IR-T1D-000004. HLA genotype labels were derived from supporting information in primary manuscripts [4,18,20] or by request to the corresponding author [21].

### Code Availability

Code used for feature discovery, model fitting, and calibration is available at github.com/kmayerb/TCR2HLA/tcrtest. A command-line python tool for running TCR2HLA with multiple cpus is available at github.com/kmayerb/TCR2HLA. An interactive serverless web-browser application is also provided: kmayerb.github.io/TCR2HLAi/. Tools for CPU or GPU-accelerated approximate TCRdist computation and network-based meta-clonotype motif discovery are available at github.com/kmayerb/tcrdistgpu.

## Contributions

Conceptualization: KMB, AFG, PB, VG, PGT

Data Curation: TMB, PR, MS, LP, DMK, KMB

Formal Analysis. KMB, AFG, AM, MP

Funding Acquisition. AFG, PGT, PB, DMK, VG, TB

Investigation. KMB, AFG

Methodology. KMB, AFG, MP, AM, PB

Software. KMB, SM, MP

Supervision. AFG, PGT, PB, VG, TMB

Validation. KMB, KV

Visualization. KMB

Writing Original Draft. KMB, AFG, AM

Writing Review & Editing. KMB, AFG, AP, EF, AM, PR, MRS, LDP, KV, SM, DMK, VG, PB, TMB, PGT

## Acknowledgements

We thank Dr. Daniel Geraghty for providing high-resolution HLA-genotyping participants in the Elyanow et al. study, Ying Lei for helpful discussion about a memory-efficient algorithm for finding mutational variants, and Dr. Liel Cohen-Lavi for helpful early discussion on HLA feature selection. The authors used an LLM (Microsoft Copilot and ChatGPT-4) to assist with proofreading and reduction in word length for sections of the manuscript text. Proofread versions of the manuscript were compared with the authors’ original text using Microsoft Word’s compare documents tool. All proposed grammatical changes were manually reviewed and accepted or rejected. All scientific content, citations, and interpretations are the authors’.

## SUPPORTING FIGURES

**Figure S1:**
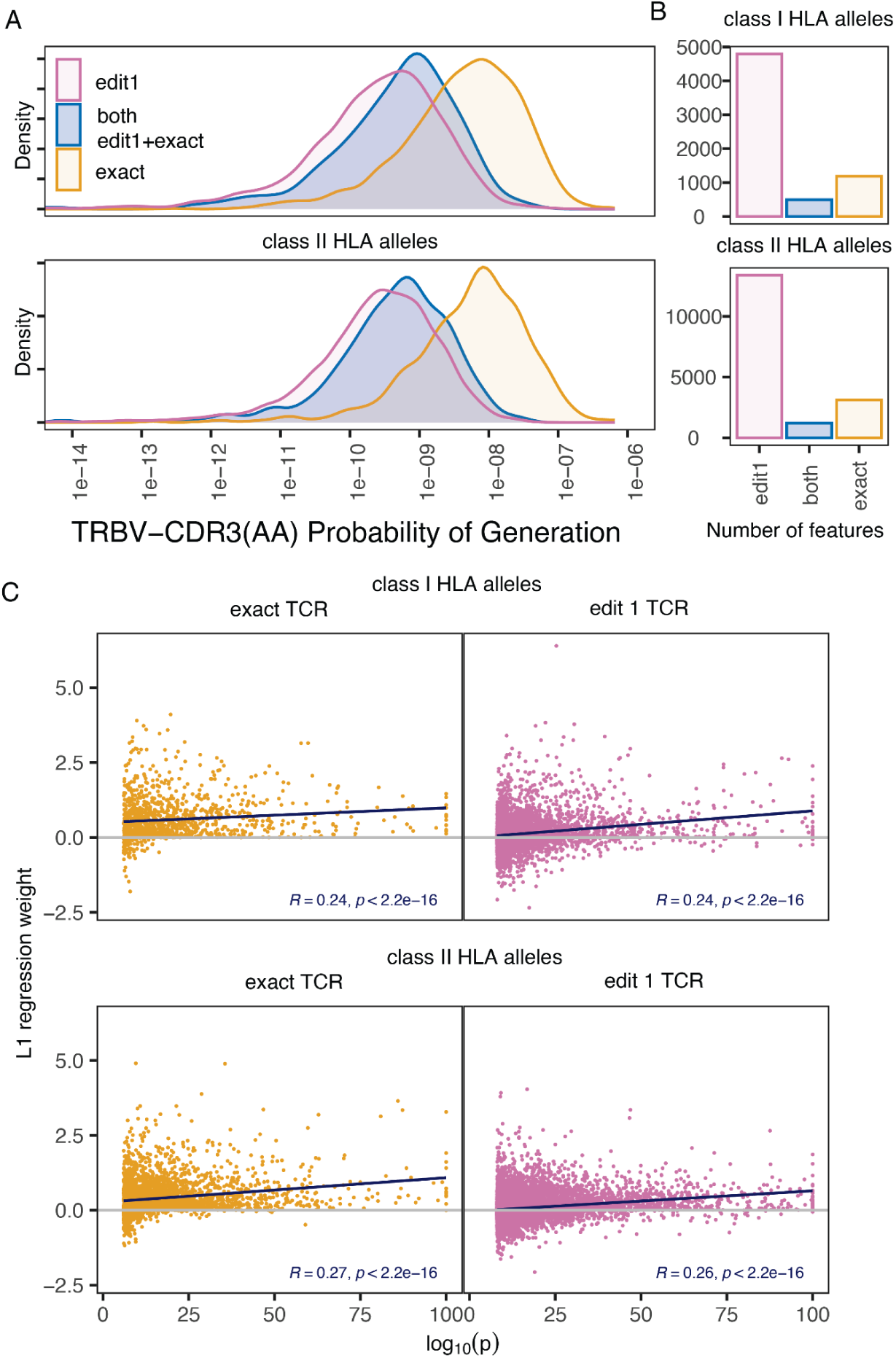
Pgen estimates and L1-regularized feature weights by TCR feature type. (**S1a**) Estimated probability of generation of TCRs (TRBV-CDR3AA) associated with common class I or class II HLA alleles. (**S1b**) Number of exact or near-exact (edit-1) HLA-associated TCR with non-zero L1-regularized regression weights. TCR features that were strongly associated with an HLA-allele by exact matching (orange), and near-exact (single mutational variant inclusive enumeration (pink), or both (blue).

**Figure S2:**
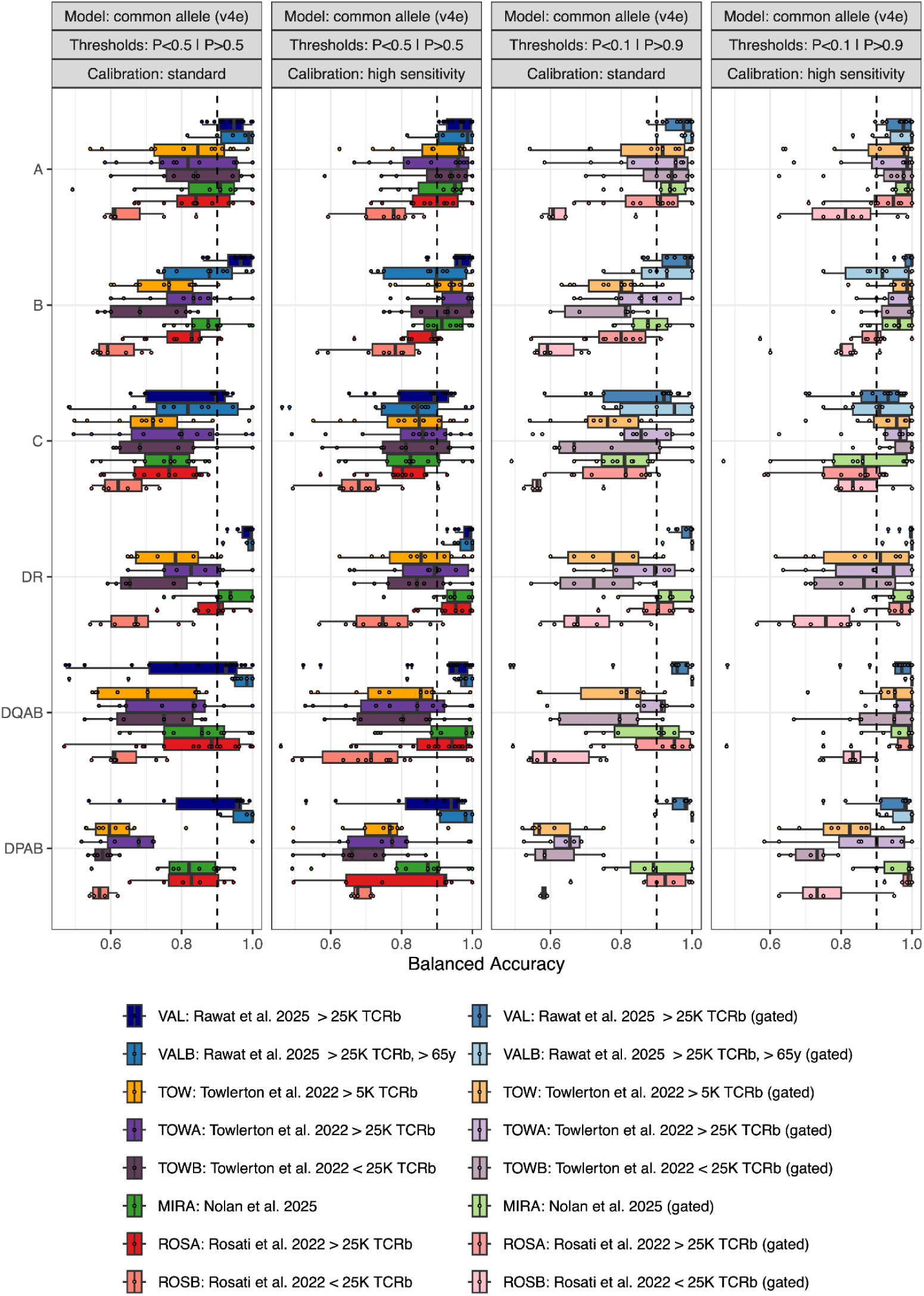
Balanced accuracy on different validation datasets. Balanced accuracy was estimated by predicting common HLA alleles across DNA-based (Rawat, Towlerton, Nolan (MIRA)) or RNA-based (Rosati) TCRseq external validation data sets. (**S2a**) Standard calibration using a P > 0.5 decision threshold. (**S2b**) High sensitivity calibration using P > 0.5 decision threshold. (**S2c**) Standard calibration with intermediate confidence predictions (0.1 > P(HLA) > .9) removed. (**S2d**) High sensitivity calibration with intermediate confidence predictions (0.1 > P(HLA) > .9) removed. The standard calibration was based on full training data. The high sensitivity calibration was based on repeated downsampling of the training data to incorporate an adjustment based on the number of unique clonotypes ranging from 5,000 to 500,000 (see Methods).

**Figure S3:**
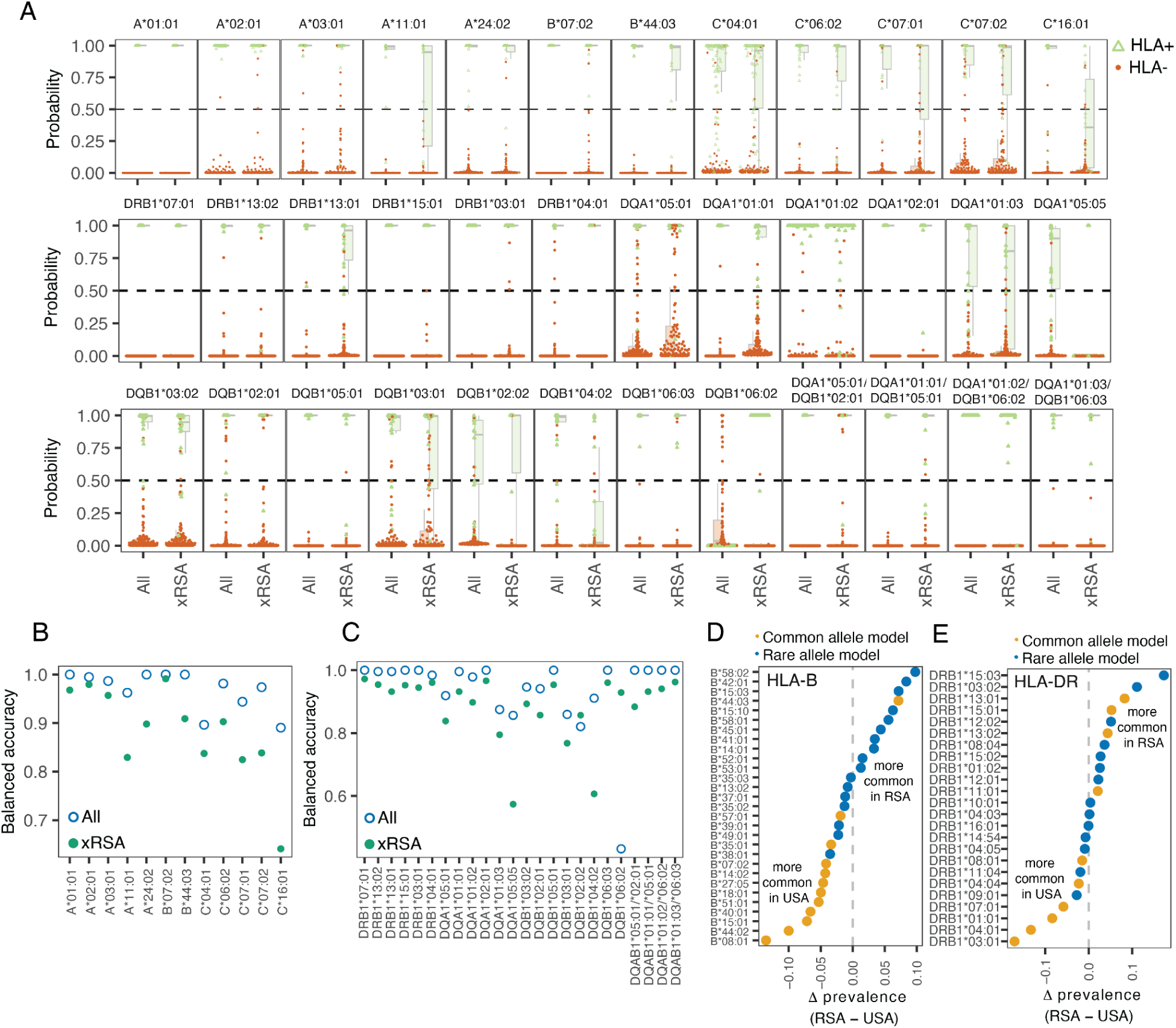
HLA-inference to geographically distinct populations. (**S2a**) For repertoires from the Republic of South Africa (RSA), calibrated probabilities were generated by models using all training data (All) or after removing RSA samples (xRSA). (**S2b,c**) Balanced accuracy of predictions of full vs. ablated models for common class I or class II HLA alleles. (**S2d,e**) Difference in allelic prevalence between the RSA and US training data for representative HLA-B (**d**) and HLA-DR (**e**).

